# Nationwide assessment of leadership development for graduate students in the agricultural plant sciences

**DOI:** 10.1101/2022.12.04.519039

**Authors:** Karen Ferreira Da Silva, Ella Burnham, Joe Louis, Douglas Golick, Sydney Everhart

**Affiliations:** Department of Plant Pathology, University of Nebraska, Lincoln, Nebraska, United States of America; Department of Statistics, University of Nebraska, Lincoln, Nebraska, United States of America; Department of Entomology, University of Nebraska, Lincoln, Nebraska, United States of America; Department of Biochemistry, University of Nebraska, Lincoln, Nebraska, United States of America; Department of Plant Science and Landscape Architecture, University of Connecticut, Storrs, Connecticut, United States of America

## Abstract

Leadership development is a universally important goal across the agricultural plant science disciplines. Although previous studies have identified a need for leadership skills, less is known about leadership skill development in graduate programs. To address this, we constructed a mixed-method study to identify the most significant graduate school leadership experiences of scientists in the agricultural plant science disciplines. The survey was deployed to 6,728 people in the U.S. and received 1,086 responses (16.1% response rate). The majority of respondents reported that they were from one of the major agricultural states and employed at one of the agricultural plant science related doctoral universities, industries, or government. Results from this survey suggest that recent graduates were more engaged in graduate school activities that offered leadership development. Key experiences in graduate school were also identified that may be used to develop future leaders. Additionally, respondents reported the greatest barrier to providing leadership development for graduate students was that it is not part of their program curriculum, however current graduate students responded differently, and identifying lack of funding to support experiences as the greatest barrier. This survey also identified the top ranked professional skills considered most important for effective leaders in agricultural plant sciences as well as respondent-driven recommendations on how graduate programs can improve leadership development. Collectively, these results can be used in the future to identify priorities for skill development and opportunities for graduate student training in leadership skills within the plant science disciplines.

## Introduction

Research shows that the most successful leaders have not only mastered technical skills, but they have also mastered professional skills [1]. Among successful business leaders and executives around the globe, talent, and leadership development programs were identified as top priority because individuals with strong leadership skills are able to ensure that projects run smoothly, and tasks are completed [2,3]. Among graduate students in the sciences, technical proficiency is considered an expectation, while those also trained in leadership skills are increasingly sought after by employers [4]. Therefore, it is critical for leadership skills to be developed in graduate students, which will make graduates more competitive for jobs and aid their long-term career development.

Leadership skills development should be incorporated into graduate programs because those with advanced degrees are often expected to lead people as part of their job duties [5]. Consequently, for individuals in advanced degree programs, access to high quality leadership training is essential to bridge the gap between leadership development and technical skills. Although educational programs in leadership exist for undergraduate students at universities across the nation, graduate programs within the science, technology, engineering, and math (STEM) disciplines typically do not provide leadership preparation. A recent survey showed that among those plant scientists who received some level of leadership training as part of their formal degree program, only 36% of scientists reported that the training received was useful for their leadership roles [6].

As the number of science and engineering doctoral degrees awarded in the U.S. has increased significantly, the proportion of graduates reporting a definite job commitment, including postdoctoral positions (postdocs), have declined in the last 10 years [7]. A survey on job availability for STEM students showed that only 13% of PhD graduates will hold an academic position in the U.S. [8], therefore requiring graduates to find jobs in non-academic roles/institutions. Indeed, an increasing number of graduate students are interested in the private sector. A survey reported that while 75% of respondents were interested in an academic career opportunity after graduation, 55% were interested in an industry position [9]. However, these graduate students are not leaving research altogether, as 80% reported the likelihood that they will pursue a research career has grown or remained unchanged since they began their Ph.D. program [9]. Hence, training programs for the scientific workforce should be tailored for these different career track options for STEM graduate students.

Graduate students in agricultural plant sciences can receive formal scientific training in a wide range of disciplines such as Agronomy, Crop Sciences, Entomology, Environmental Sciences, Horticulture, and Plant Pathology. These students are often advancing their technical knowledge to address direct applied issues to the society. The significant increasing demand for food production from the global population is projected to rapidly grow from 7.7 billion in 2019 to approximately 11.2 billion by 2100 [10], which underscores the need to offer professional skill training for agricultural plant scientists beyond technical training. Such skills, including leadership, are considered value-added skills to encourage future scientists to effectively tackle these future challenges and demands. Leadership skills when added to adequate technical competencies can turn these professionals into highly competitive candidates in the marketplace as they are perceived with potential talents and capabilities to address the gap of food security and agricultural production needs.

Although leadership is an important employability trait, employers are concerned that applicants are deficient in the core, transferable skills that they will need to succeed [11]. Employers report that 30% of applicants were unable to demonstrate leadership experience or skills. Educators surveyed in this study also reported that leadership is a skill that students are least likely to be equipped with when they enter the job market. A recent survey also found that employers rated new graduates to be the least proficient in leadership competencies compared to all other job ready skills [12]. Remarkably, another study showed recent graduates over-estimate their professional skill proficiency compared to employer ratings of their proficiency [13]. Among professional skills, leadership was ranked in the top three, which also had the greatest disparity in proficiency, wherein 70.5% of the students considered themselves proficient in leadership skills, but only 33% of the employers agreed.

To better understand the needs, experiences, and limitations for leadership training, we developed and implemented a survey to assess leadership development experiences and needs for future agricultural plant scientists nationwide. While substantial differences may exist in professional needs and training across STEM disciplines, a focus on the agricultural plant sciences minimizes such differences to enable a concise investigation. Specifically, this study aimed to identify gaps and opportunities in leadership training for universities to improve professional development opportunities for graduate students focused on plant science related disciplines and provide more information to prepare the next generation of leaders in science.

Specifically, the goals of this study were to:

1. Identify graduate student experiences and activities that career professionals feel significantly assisted their leadership development;
2. Determine key professional skills needed among scientists to be effective leaders;
3. Identify potential barriers to providing leadership development in graduate school;
4. Collect and synthesize suggestions for graduate programs should use to increase leadership development among plant scientists; and
5. Determine if there are differences in experiences and leadership skills needed for industry and academic career tracks.

## Materials and Methods

### Survey construction

A web-based survey was designed with seven blocks of questions, totaling 50 questions. The survey was developed by the authors based on previous leadership surveys [6,14]. The questions were constructed with the goals of addressing the study goals. Prior to deployment of the nationwide survey, interviews were conducted with university leaders and plant science industry professionals to evaluate the initial survey questions. The survey went through two rounds of post-interview revisions.

Questions about professional skills were created following the organizational structure already established by a previous study that focused on STEM students and employability skills [14]. In that study, the authors aggregated the list of skills from a comprehensive examination of over 80 articles and publications on employability skills. Further, the authors identified a collection of skills that were associated with effective leadership and were directly related to effective leadership preparation. In the present study, we used the same collection of skills and these were presented to the participants to be able to rank the three most important skills based on their own leadership experience.

Some questions allowed for participants to provide a short-written answer for their opinion and recommendation in leadership development. These open-ended questions were designed to provide the participants the opportunity to elaborate about their experience on leadership and to gather more information on the wide array of activities that these professionals are being exposed to in their current employment setting.

### Pilot testing and survey validation

As part of the survey development process, the survey questions underwent testing to see if the measures and contents provide the expected outcome. First an internal survey content validation was performed by using literature review and assessment by the authors to ensure that items deemed essential were maintained and to eliminate undesirable items [15]. Further, we conducted pilot testing of the survey with eight participants that have leadership experience, received or were pursuing Ph.D. degree in the agricultural plant sciences field, and were working in either academic or industry employment settings. These participants were asked to take a preliminary version of the survey that had both qualitative and quantitative questions, following which, an in-person interview was conducted using semi-structured questions formatted to learn more about their leadership experience and to collect their feedback about the pilot survey. The authors discussed the survey and interviews responses and questions were revised based on their feedback.

The survey was designed for respondents to complete during a two-week period. An extended completion window was allowed due to large number of questions and the fact that questions had many response options. Surveys were deemed a desirable data collection method as we wanted to best assess characteristics of a wide variety of professionals in the agricultural plant science fields. Additionally, some questions like those self-evaluating abilities may be more stigmatizing or difficult for people to express honestly to an interviewer in a face-to-face setting. Self-administered survey with standardized questions can be more reliable compared to qualitative methods, since the investigators are not able to influence responses when compared to a face-to-face interview [16]. The web-based questionnaire was hosted on Qualtrics Survey Software [17].

### The Survey

Standard demographic questions used in the U.S. census (e.g., gender, age, ethnicity, and race) were included. Other questions related to their educational background and degree(s), the types of activities respondents had during these degrees, job position description, employment type, and location where the respondents work or study (state within the U.S.). Additionally, information about the size of their current organization of employment, and questions related to their experiences with leadership and management were surveyed. Questions regarding leadership experience, graduate program training experiences and activities, barriers on leadership development, professional skills, and open-ended questions to capture recommendations and opinions from participants were also included.

The first set of questions were used to identify current professionals in the workforce with graduate degree who possess high level managerial experiences and also to capture current graduate students and professionals who possess a graduate degree and have or had a formal or informal leadership experience. The survey defined *formal leadership* as an appointed or elected position or role in a professional or non-professional setting and *informal leadership* as tasks performed in an unrecognized (informal) position, with individuals or groups. These definitions were provided to assist participants in identification of roles that may involve leadership skills, not just those with formal (appointed or elected) supervisory responsibilities.

To gain further insight about their leadership experiences, we asked participants to list other types of professional involvement, in addition to their research and teaching responsibilities, such as participation in professional societies, internships, and community services. To gauge the impact of leadership experiences, participants were also asked to rate the importance of their experiences and activities for their professional and leadership success.

Lastly, a list of experiences and activities pursued in graduate school and a list of potential barriers in leadership development was provided with the goal to identify opportunities to improve leadership development for current and future graduate students. Respondents were asked to rate the importance of such individual experiences and also their perception of suggested barrier to offering leadership development training in the academic setting.

All the investigators of this study completed the Collaborative Institutional Training Initiative (CITI) certification required by the University of Nebraska-Lincoln’s (UNL) Institutional Review Board (IRB) and were maintained during the life of the project. This project has been certified by the IRB as exempt, category 2, project number: 20190118764EX.

### Recruitment

The purpose of this study was to gain insight into the importance and role of various experiences in agricultural plant science graduate programs on leadership development relevant to professional careers. The survey was sent out to graduate students and professionals from agricultural plant sciences field. Therefore, the researchers purposively targeted recruitment of members of the American Society of Agronomy, the Crop Science Society of America, and the Soil Science Society of America, Entomological Society of America, and the American Phytopathological Society. These organizations have the highest concentration of members who are students or career professionals in the agricultural plant sciences, which includes: entomology, plant pathology, agronomy, horticulture, soil science, environmental sciences, and related disciplines.

The recruitment process attempted to provide a representative subsample of each society’s membership, since collecting contact information of every member would provide more contacts than needed for the purpose of this study. Participants were haphazardly selected by scrolling through the online membership list provided by each society. This recruitment methodology did not intend to provide an equal chance of selection between demographic or other groups (e.g., equal ratio of males and females or industry vs. academic employer), rather it was designed to create equal number of participants from each society.

A total of 6,728 email contacts were collected and used for survey distribution through Qualtrics Survey Software. Qualtrics offers an online survey software service which allows researchers to create, customize, and modify research projects [17]. This software has an automated feature that generated the links that were sent simultaneously to all participants. A letter regarding the purpose of the study and research participant consent was included with the survey link.

### Distribution

Surveys were distributed via email in January of 2019. The email contained a message with a notification about their selection and a link. After the participants received the first email containing the link to access the survey, follow-up emails were generated by Qualtrics each week as reminders to complete the survey. The follow-up emails were used since previous studies have shown that sending follow-up messages increased the percentage of people who return their questionnaires [18]. All responses were anonymous and no personal information was collected during the survey.

### Data analysis

#### Data validation and filtering, and response partitioning

Data validation ensures that the survey questionnaires are completed and represent accurate response data [19]. The data file was downloaded in Microsoft Excel format and kept in a password protected file. From there, participants, whose questions with relevant variables used for data analysis were not answered, were removed from the analysis. Such variables that eliminated a response from analysis were mainly those related to the demographic portion of the survey, which were used to classify participants’ professional profile and leadership training. Blank responses were removed from analysis.

#### Data coding and statistical analysis

Survey question data consisted of a combination of nominal, ordinal, interval, and ratio measurements. The type of measurement variable determined the type of statistics and data analysis that were applied determined the type of information obtained from the survey [16]. Analysis of the quantitative data and graphs and charts were constructed in R software [20] using the following packages: naniar [21], gplots [22], reshape2 [23], ggplot2 [24], readr [25], tidyverse [26], dplyr [27], likert [28], nnet [29], HH [30], Hmisc [31], maps [32], mapdata [33], maptools [34], rgeos [35], rgdal [36], mapproj [37].

#### Thematic analysis

Written answers were initially themed and coded by the authors. Categories were further examined and coded by the UNL Bureau of Sociological Research (BOSR) staff member with expertise in qualitative data analysis and coded into common themes. The data consisted of respondents that identified as professional in either an academic or industry employment setting. There were 13 common themes in total, with an extra category that was designated for missing data. Each answer was coded into at least one theme but included the possibility of being coded into multiple themes. After each answer was coded into at least one of the common themes, they were further coded into subcategories to facilitate analysis.

## Results

### Demographics

Of the 1,086 participants that completed the survey, 835 participants, including both professionals and graduate students, were selected for further data analysis after data validation and filtering. The distribution of participants by geographic location in the U.S. (Fig. 1A), employment type (Fig. 1B), and age versus years after graduation (Fig. 1C) is illustrated in Fig 1. The U.S. map shows the widespread distribution of respondents in each state. Geographically, the states with the largest number of respondents were California (9.6%), North Dakota (6.1%), and Florida (5.3%), and the only state without respondents was Alaska. The distribution of participants by employment type showed most were members of doctoral universities (46.9%), followed by industry (18.3%), and government (17.9%). There was a positive correlation between years of birth and graduation, which enabled to categorize the post graduate career stages into groups by years since graduation as of ‘Early-career’, ‘Mid-career’, and ‘Late-career’. For questions related to professional experiences, only respondents that identified as being part of the workforce (n = 674) were analyzed.

**Fig 1.**
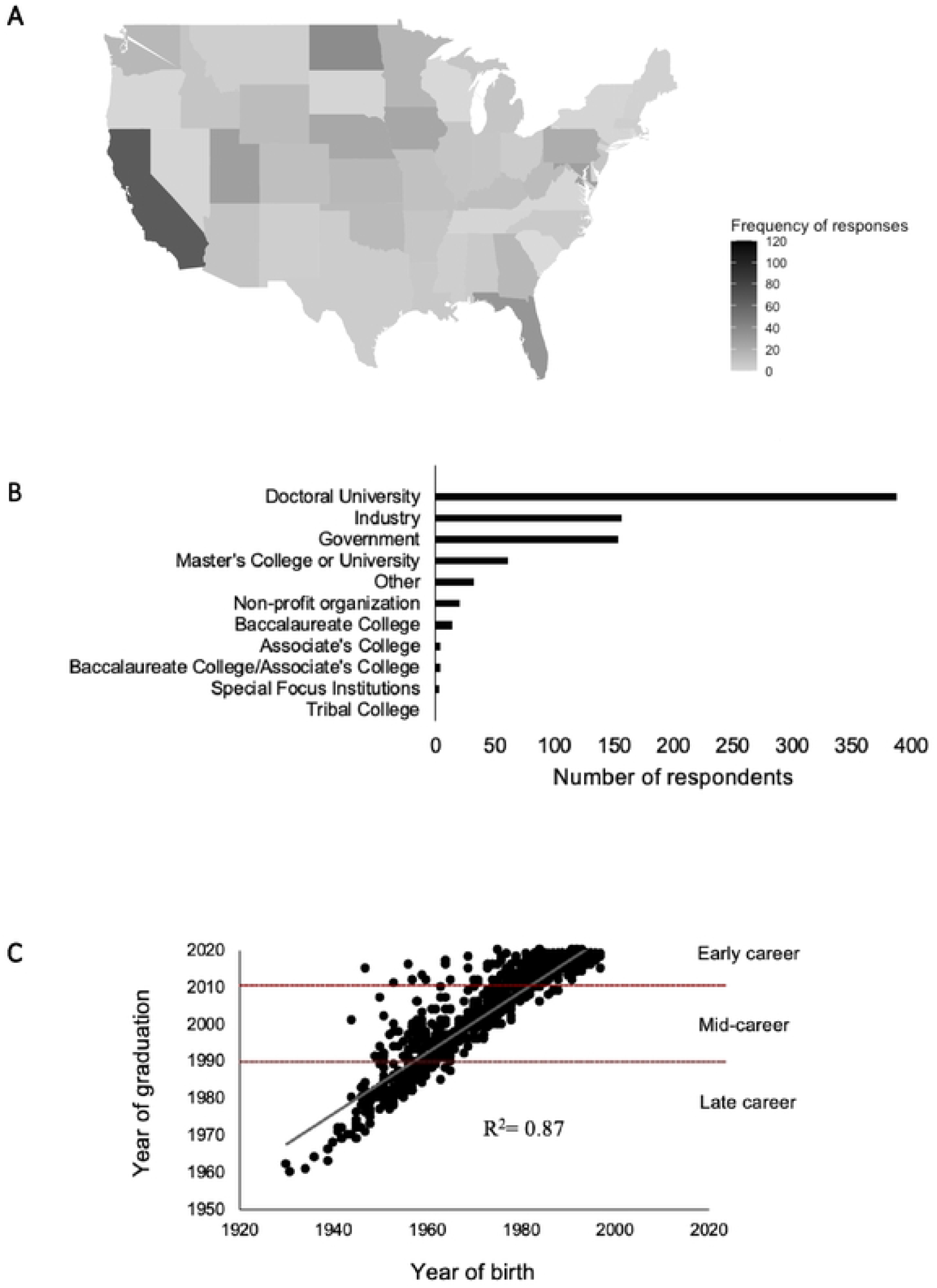
Demographics of the respondents participated in the survey. (A) Employment type distribution: Number of respondents to the survey, including both professionals and graduate students. (B) U.S. Geographic distribution: The U.S. map shows the widespread distribution of respondents in each state (not including Alaska, Hawaii, and Puerto Rico). (C) Birth year versus Graduation Year: The year of graduation and birth of respondents was correlated.

### Significance of leadership training in graduate programs

When asked about the importance of their own graduate programs in preparing students for leadership roles in the future professional career, more than half respondents who received their graduate degree (n = 668), indicated that graduate programs were significant (34.3%) or very significant (21.3%), compared to those that indicated they were not significant (34.1%). Similarly, among enrolled graduate students (n = 160), more than half perceived graduate school as significant (32.5%) or very significant (24.4%) in preparing for future leadership roles and, with fewer (39.4%) that perceived graduate programs as a non-significant role in preparing them for future leadership roles.

When looking at industry versus academic employment type, academics were slightly more positive when rating the importance of graduate programs to their career success as leaders. Among academics respondents (n = 267), 37.1% and 21.0% rated their graduate program as significant and very significant for their success as leader, respectively, whereas industry respondents (n = 153) 33.3% and 17.0% rated their graduate program significant or very significant for their success as leaders.

### Professional Activities in Graduate School

Survey participants were asked to share the types of professional activities they performed as a graduate student. The following type of activities were given to them to choose among all those applied during their graduate training: ‘Research’, ‘Teaching’, ‘Lecturer (unsupervised course instruction)’, ‘Outreach’, ‘Extension program development/ delivery’, and ‘Service (e.g., committees, clubs, governance)’. To investigate whether the types of activities available to graduate students has changed during the past few decades, we categorized responses according to the number of years since graduation as illustrated in Fig. 2.

**Fig 2.**
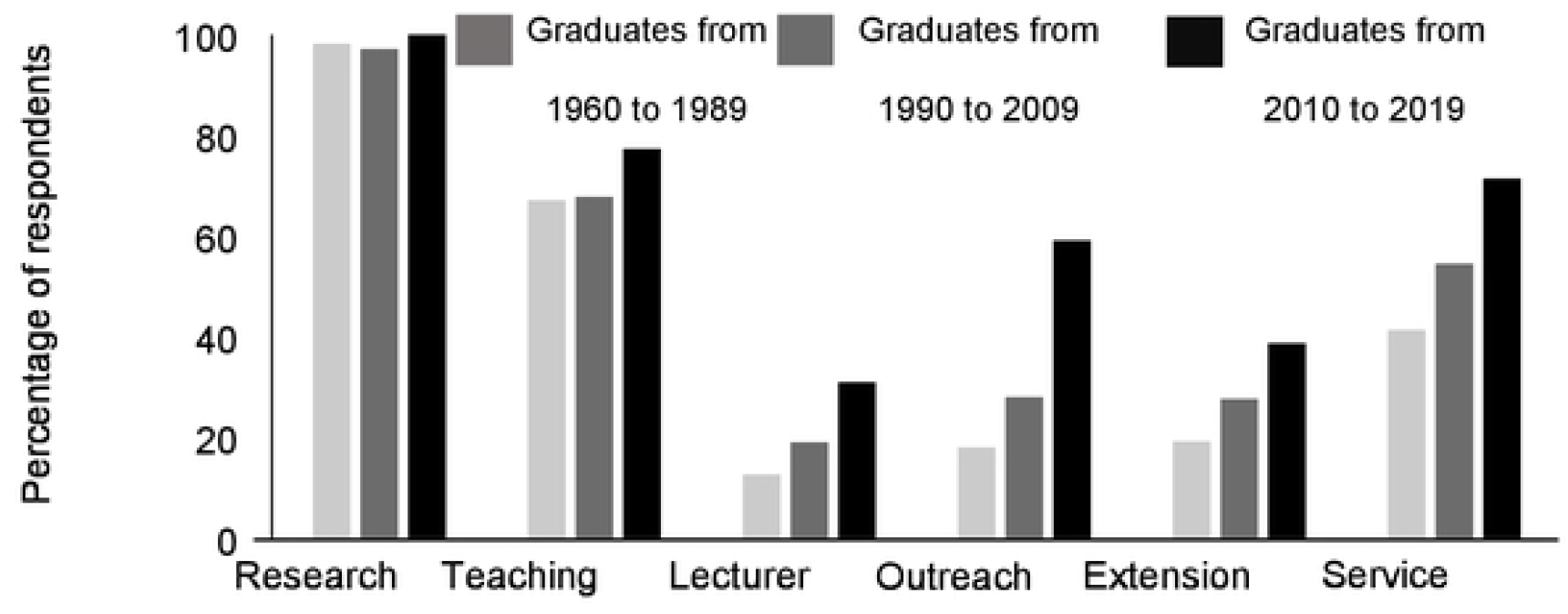
Number of activities in graduate school. A total of 665 participants responded that they participated in one activity or all of them. For Research (n = 656), TA (n = 471), Lecturer (n = 142), Outreach (n = 238), Extension program development/delivery (n = 194), Service (e.g. committees, clubs, governance) (n= =378). Graduates from 1960 to 1989 (n = 167), Graduates from 1990 to 2009 (n = 279) and graduates from 2010-2019 (n = 219).

Overall, there was an increase in the percentage of respondents who were recent graduates (from the last decade), in each activity type performed during graduate school. Greater variation was observed for outreach, in which a 31.1% increase in respondents reporting outreach experience was observed when comparing recent graduates to those graduating from 1990 to 2009. A similar trend was observed when comparing the number of activities per respondent, which showed recent graduates reported the highest number of activities per respondent (3.75 ± 1.45 activities per respondent), while graduates from 1990-2009 and 1960-1989 reported 2.91 ± 1.46 and 2.52 ± 1.33 activities per respondent, respectively. Therefore, the investigators were interested in understanding if there was a correlation between the number of activities and professional leadership roles. To answer this question, respondents that self-identified as holding a management role were selected for further analysis. A summary of the mean number of graduate school activities by level of management is shown in Table 1. On average, upper managers had 2.76 ± 1.57 activities in graduate school, middle managers had 3.02 ± 1.49 activities in graduate school, and lower managers had experienced 3.24 ± 1.46 activities in graduate school.

**Table 1.** Management type versus number of activities with median, mean and standard deviation.

|  | Sample Size | Min | Q<sub>1</sub> | Median | Mean | Q<sub>3</sub> | Max | SD |
| --- | --- | --- | --- | --- | --- | --- | --- | --- |
| Lower | 33 | 1 | 2 | 3 | 3.24 | 4 | 6 | 1.46 |
| Middle | 127 | 0 | 2 | 3 | 3.02 | 4 | 6 | 1.49 |
| Upper | 83 | 1 | 1 | 2 | 2.76 | 4 | 6 | 1.57 |

However, a two-sample independent t-test was conducted and showed there was a significantly greater number of activities reported by lower managers than both middle and upper-managers (P = 0.017, t = 2.124, and df = 241, Table 2). Examining this further showed that the top three activities reported, regardless of current management roles, were ‘Research’, ‘Teaching Assistant’, and ‘Service (e.g. committees, clubs, governance)’.

**Table 2.** Frequency of respondents in management roles who performed activities during graduate school.

|  | Research | Teaching Assistant | Lecturer<sup>a</sup> | Outreach | Extension<sup>b</sup> | Service<sup>c</sup> |
| --- | --- | --- | --- | --- | --- | --- |
| Lower | 1 | 0.82 | 0.15 | 0.33 | 0.27 | 0.67 |
| Middle | 0.97 | 0.68 | 0.27 | 0.28 | 0.28 | 0.53 |
| Upper | 0.98 | 0.59 | 0.20 | 0.30 | 0.25 | 0.43 |

<sup>a</sup>Lecturer is unsupervised course instruction; <sup>b</sup>Extension is the development and delivery of extension programs; <sup>c</sup>Service can be participation in committees, clubs, and governance.

### Experiences

Participants that reported to have either formal or informal leadership experience were asked to rate the overall importance of having a related professional experience during graduate school to their current leadership success. Results showed the majority of respondents (42.1%) rated ‘Observing other leaders’ as the most significant experience, followed by ‘Job experiences’ (40.8%), and ‘Mentoring students in field or lab research’ (33.1%). Interestingly, even though ‘International study/experience’ presented a lower number of respondents (n = 265), its importance was shown to be high (41.1%). On the opposite side, ‘Formal college course that taught leadership skills’ and ‘Engaging in campus or inter-institutional leadership preparation programs, e.g., Center for Integration of Research, Teaching, and Learning (CIRTL), preparing future faculty’ were generally rated as non-significant experience (54.3% and 47.7%, respectively). Among experiences listed in the ‘Other/not listed’ category included: ‘self-drive’, ‘LEAD21 program’, ‘understanding different cultures and working with international research’, ‘church programs’, ‘business courses and MBA program’, ‘field work’, ‘Toastmasters’, ‘serving on department/university committees’, and ‘participation in civic organizations’.

When comparing responses from participants in industry to those in academia we found no significant differences (*P* > 0.01 for each and df = 2); however, some trends were observed. Experiences such as ‘Formal college courses’, ‘Engaging in campus or inter-institutional leadership preparation programs, e.g., CIRTL, preparing future faculty)’, and ‘Mentorship from the graduate advisor’ were more highly rated among academic employees, whereas ‘Internships’ and ‘Job experiences’ were more highly rated by industry employees.

In the analysis, we were also interested to evaluate if the number of years in leadership experience was related to leadership success. Responses from participants with formal leadership experience (n= 626) were quantified and ranked according to the experiences that were most valuable for their leadership success in Fig. 3.

**Fig 3.**
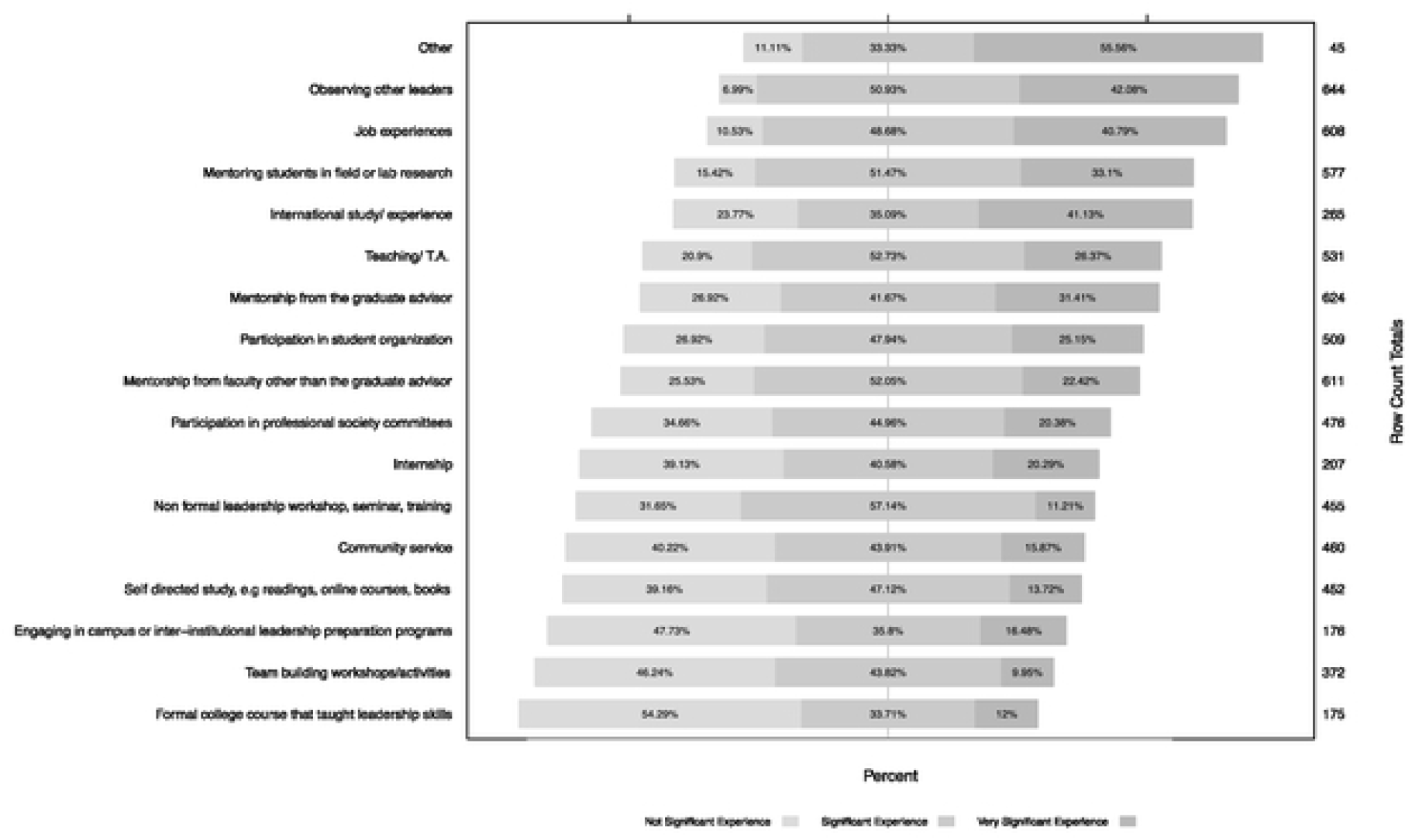
Experiences in graduate school in the order of importance for leadership development. Comparison of 17 graduate program experiences rated as not significant, significant, and very significant. Row count totals represent the number of respondents that rated each experience.

### Barriers

All participants were asked to rate the overall importance of a prepared list of situations that could pose as a barrier to leadership development of students during graduate school (Fig. 4) and results showed these were significantly different (*P* < 0.001).

**Fig. 4.**
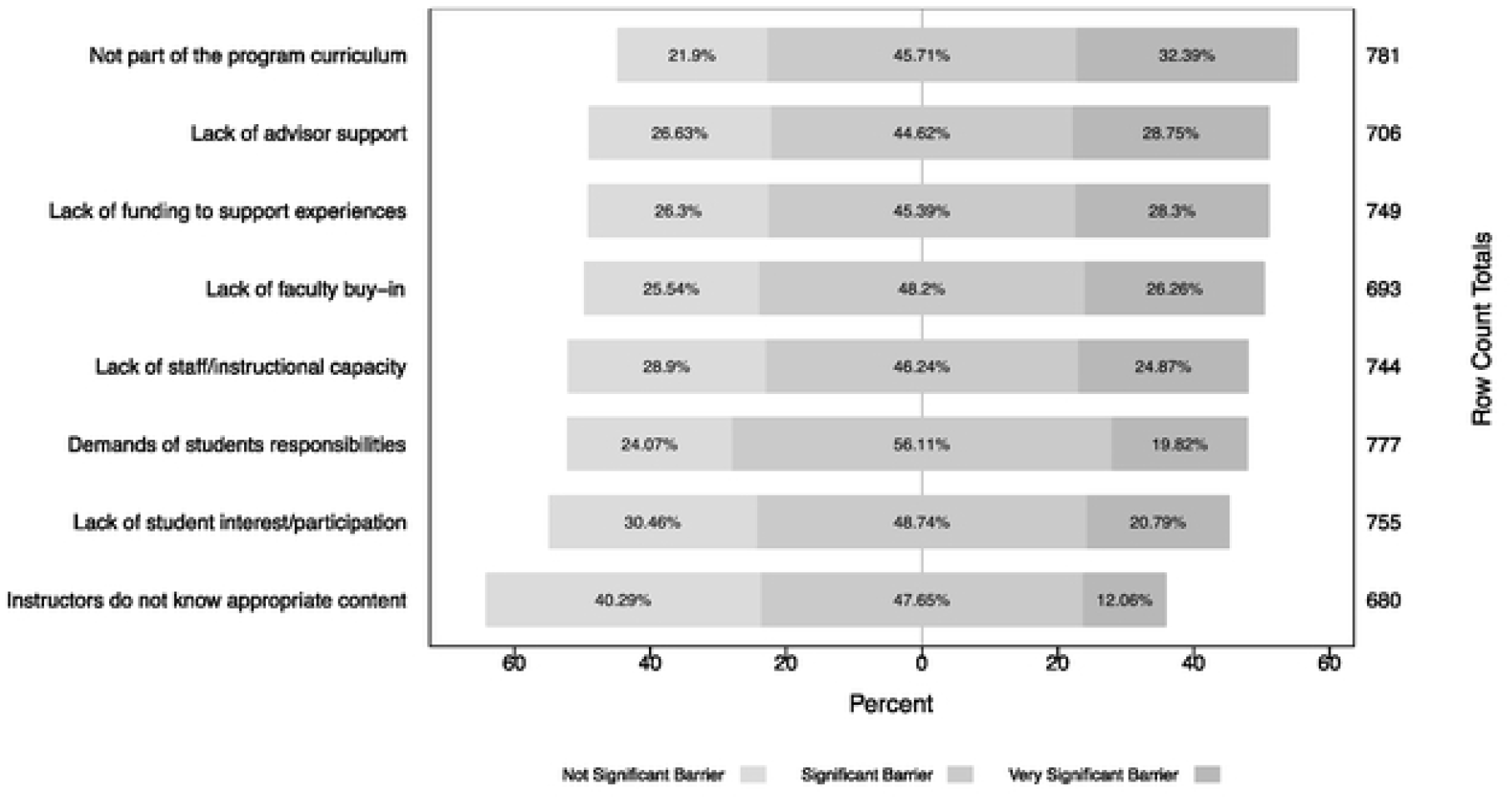
Barriers for making formal and informal leadership experiences available to graduate students. Comparison of eight examples of possible barriers rated as not significant, significant, and very significant. Row count totals represent the number of respondents that rated each experience.

Among respondents (n = 824), ‘not being part of the program curriculum’ (32.4%), followed by ‘students lack advisor’s support’ (28.7%), and ‘lack of funding to support experiences’ (28.3%), were considered the top three most significant barriers in offering leadership development in universities. Comparing responses between those who have received their graduate degree (n = 664) and current graduate students (n = 160) showed no significant difference in the importance of these barriers (*P* > 0.01; df = 2). However, some differences were observed in the rank order of barriers between groups. Among respondents who received their graduate degree, ‘not being part of the program curriculum’ (30.7%), followed by ‘lack of funding to support experiences’ (23.5%), and ‘demands of students responsibilities’ (23.5%) were perceived to be the top 3 most significant barriers in offering leadership development in universities. Among current graduate students, ‘lack of funding to support experiences’ was the most significant (35.6%), followed by ‘not being part of the program curriculum’ (30.8%) and ‘students lack advisor’s support’ (30.6%).

### Skills

Respondents were asked to select and rank the top three skills that in their opinion were considered the most important for an effective leader. Table 3 represents the summary of the top three most important skills for a successful leader ranked among professionals (n = 674). The most important skills for an effective leader were identified as: ‘listen effectively’, ‘problem-solving’, ‘well-developed ethic, integrity and sense of loyalty’, ‘collaborative’, and ‘being strategic and seeing the big picture’.

**Table 3.**
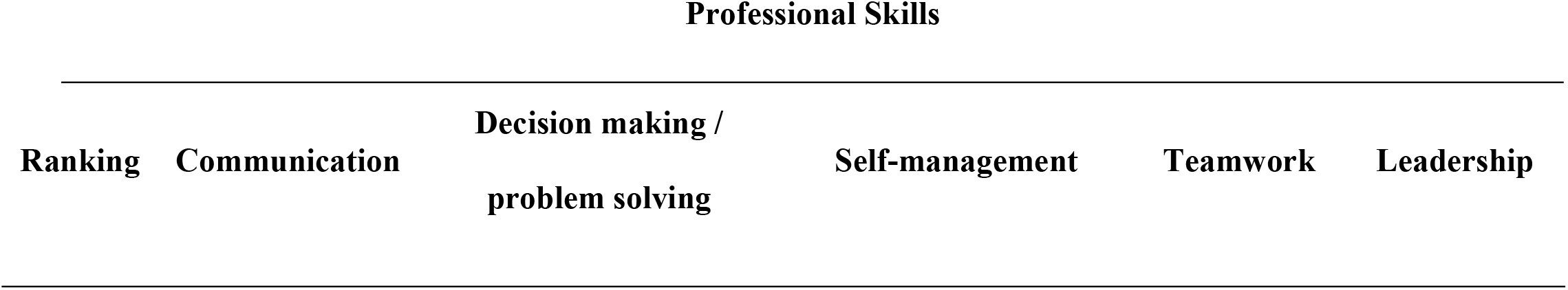
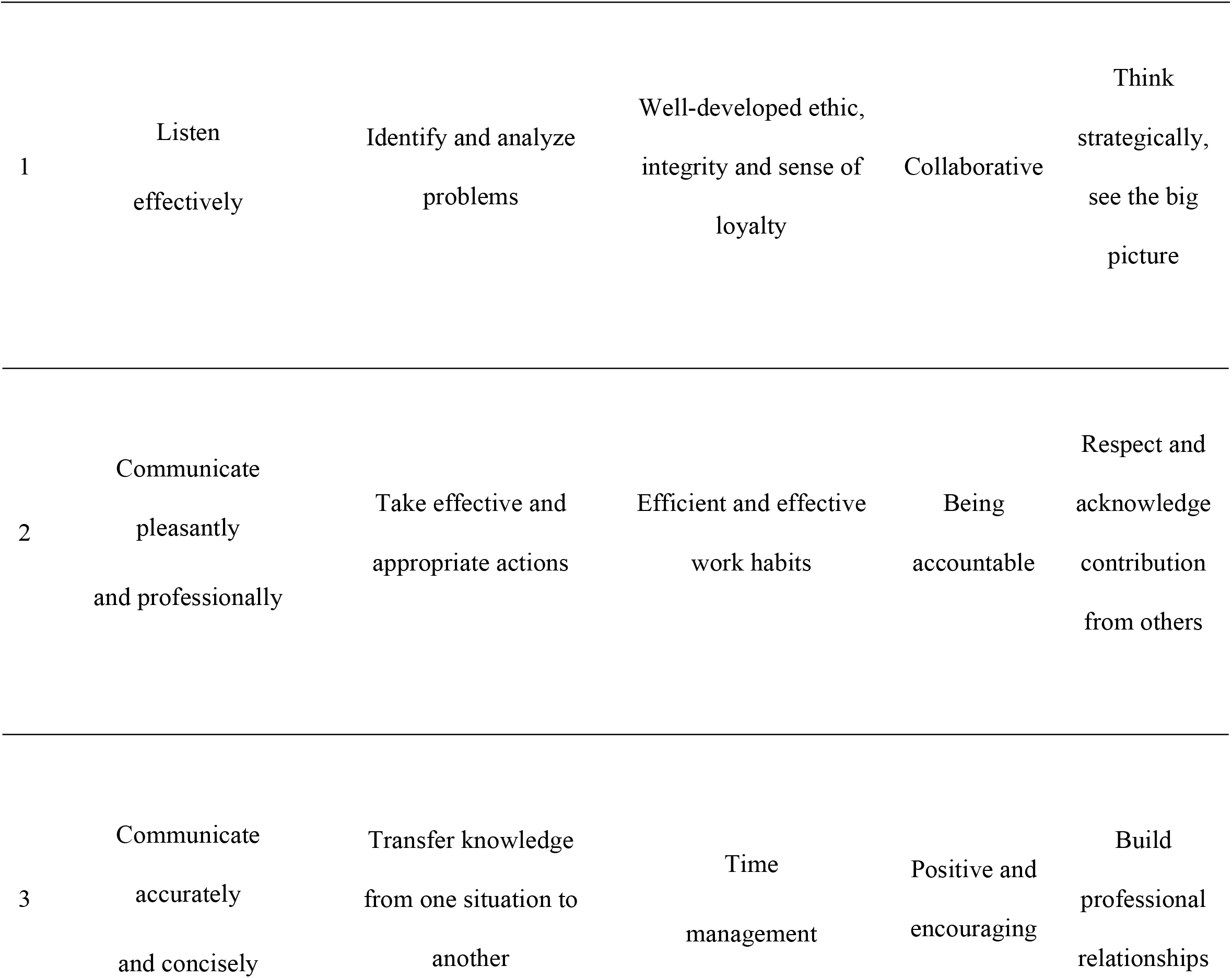
Top ranked professional skills among agricultural plant science professionals. Five skill clusters are represented in the first row followed by the top three most ranked skills by survey respondents in professional roles.

| Professional Skills |  |  |  |  |  |
| --- | --- | --- | --- | --- | --- |
| Ranking | Communication | Decision making /<br>problem solving | Self-management | Teamwork | Leadership |
| 1 | Listen<br>effectively | Identify and analyze<br>problems | Well-developed ethic,<br>integrity and sense of<br>loyalty | Collaborative | Think<br>strategically,<br>see the big<br>picture |
| 2 | Communicate<br>pleasantly<br>and professionally | Take effective and<br>appropriate actions | Efficient and effective<br>work habits | Being<br>accountable | Respect and<br>acknowledge<br>contribution<br>from others |
| 3 | Communicate<br>accurately<br>and concisely | Transfer knowledge<br>from one situation to<br>another | Time<br>management | Positive and<br>encouraging | Build<br>professional<br>relationships |

### Thematic analysis

All respondents were asked to elaborate on how graduate programs can better prepare students for leadership roles and leadership development in their current organization. Data consisting of respondents from industry and academia (n = 425) were divided into 13 main themes. The main themes were: ‘opportunities’, ‘formal programs’, ‘communication skills’, ‘teamwork’, ‘mentor students’, ‘mentor undergraduate students’, ‘self-development’, ‘business’, ‘encouragement’, ‘real word experience’, ‘education’, ‘professional’, and ‘other’. Themes and their definitions were defined based on patterns observed in the text responses and are illustrated on S1 Table.

**S1 Table**. Themes and their definitions developed based on patterns observed in the text responses.

Overall, the top recommendations for improving leadership were to ‘provide broad opportunities to develop leadership’ and ‘formal programs with formalized training’. Respondents recommended a couple of options within each theme to increase opportunities for students for leadership roles and leadership development that would propose that the student to be pushed “out of their comfort zone” as one respondent remarked. Responses were focused in organizing scientific events and participating in committees and student organization, being involved in administrative roles, participating in recruitment and onboarding training for new hires, understanding grant budgets, international research opportunities, and field courses. For formal programs, respondents provided examples that would give structured formal training such as workshops, courses, being part of the curriculum or a capstone course, seminars, active training sessions, training programs offered by faculty, and graduate certificates.

Real world experiences were mentioned as linking students to real life situations with practical experiences. Examples included participating in off campus organizations, exposure to non-academic careers such as industry jobs, internships, job shadowing, connecting with businesses, and interacting with the stakeholders, costumers, constituents. One participant responded “Exposure to the private and public business sector (…) through internships or bringing in companies to talk to students as they go through their studies. Attendance at industry-related conferences and meetings (…) Field Days and Grower Meetings. Anything that would get the student out of the classroom to visit the reality of the career they are studying.”

Another recommendation that was provided is related to mentorship. Mentorship could originate from the student’s advisor, faculty, and/or administrators to provide oversight and guidance. One respondent mentioned: “I think the single most important aspect of leadership development (…) is the ability to identify and recruit informal mentors who are experienced, strategic, and are not involved in your formal evaluation in any way (…) but I believe it should be encouraged by graduate advisers and has been instrumental to my success at every stage of my career from undergrad to present (…)”. Mentoring and leading by example was another comment identified in the survey.

When comparing industry versus academic employment settings, there were a couple of differences in the order of importance for these recommendations. For example, real world experience was chosen at much higher percentage in industry (24.2%) than for academics (8.2%). Another significant difference was observed for communication skills that was more frequently selected by industry (21.8%) when compared to academics (7.3%). The importance of teamwork was also different. Activities that participants recommend to improve written and oral communications were: engaging social activities, public speaking and communication class, communicate science through presentations to a diverse audience, communicating problems and progress. Conversely, academics more frequently rated ‘encouragement’ and ‘professional development’ (15.9% and 14.5%, respectively) compared to participants from industry (8.1% and 2.4%, respectively). Encouragement of students to pursue leadership development and roles, and professional development was shown to be important. When mentioned, encouragement and support was proposed to come from main advisor, and departmental unit chair/director. The full list of themes is represented in Table 5.

**Table 5.** Recommendation for leadership development in academia and industry.

| Theme | Overall (%)<br>n = 331 | Industry (%)<br>n = 124 | Academia (%)<br>n = 207 |
| --- | --- | --- | --- |
| Opportunities | 33.84 | 25.00 | 38.65 |
| Formal programs | 33.84 | 30.65 | 35.75 |
| Real world experience | 14.20 | 24.19 | 8.21 |
| Mentor students | 14.20 | 14.52 | 14.01 |
| Communication | 12.99 | 21.77 | 7.25 |
| Encouragement | 12.99 | 8.06 | 15.94 |
| Other | 12.69 | 15.32 | 11.11 |
| Teamwork | 10.88 | 14.52 | 8.21 |
| Professional | 9.37 | 2.42 | 14.49 |
| Self-development | 9.37 | 11.29 | 8.21 |
| Mentor undergraduates | 6.65 | 3.23 | 8.70 |
| Education | 6.34 | 6.45 | 6.28 |
| Business | 5.14 | 8.87 | 2.90 |

Another pattern identified by theming analysis was categorized as ‘leadership skills’; however, the results of this theme are not reported here because these responses contained seemingly unrelated examples. We speculate this happened largely because the term ‘leadership’ was the subject of many answers as they described the relationship of various activities to leadership development. Interestingly, however, this theme did allow us to identify that there were a small minority of respondents that did not see the need to leadership skill development during graduate school. Those respondents described leadership development as being outside the scope of the graduate student’s scientific training process which has a primary objective to develop scientists (i.e. technical skills training), and that leadership development activities will occur after graduate school. For example, one respondent answer is quoted “(…) graduate programs in plant sciences and agricultural related fields should focus 95% of their attention on the formal science education. Leadership training and opportunities will present themselves to leaders over time (…).”

## Discussion

Overall, this survey yielded a higher than expected response rate of 16.1%, whereas a 10-15% is considered the ideal response rate for external surveys [38]. This participation rate could be reflective of increased general interest in and need for leadership development for future plant scientists. Although this topic may be of direct relevance and interest to graduate program administrators, there are no standardized methods to easily and uniformly collect information from graduates after they have left their educational institutions [39], making it more challenging to collect such data for the purpose of improving the quality of graduate programs. Thus, data collected and summarized herein represent a valuable resource and will serve to provide contemporary insight into the need for leadership training within graduate programs in the plant sciences.

This survey was intended to improve our understanding of how particular activities from graduate programs were relevant for career professionals who have graduate degree in the plant sciences. As formal leadership positions often include a responsibility for managing or supervising employees, we used that trait as a means to filter responses from our survey and identify activities from graduate school that could be linked to a successful transition into leadership roles. We did not find evidence in our data to support that an increasing number of activities in graduate school was associated with a particular level of management or professional role of the respondent. One possible explanation for this is that the graduate student expectations and experiences have changed over time, resulting in large differences between early- and late-career professionals. This change was observed in Fig. 2, where more recent graduates (2010-2019) reported a higher average number of activities in nearly all categories compared to those that graduated in earlier years, aside from the most traditional categories (research and teaching). Nonetheless, service and outreach experiences as graduate students were frequently reported among those reporting to hold a managerial role. In our study, examples of service were provided to the participants and included, for example, participating in committees, student government, and outreach in the community. Such activities involve more social skills than the most technical focused activities in higher education (e.g., teaching and scientific research), which demand more analytical skills. This shift in engagement level could be related to the increasing perception of the need to integrate agricultural sciences with other activities related to societal needs [40]. The pool of potential student candidates in the agricultural disciplines is becoming more diverse and there is now a place for students who are interested not only in scientific aspect of food production and agriculture, but also in business, economic, environmental, and social issues. Additionally, service participation indicates positive effects on academic performance, values, self-efficacy, and leadership [41]. As a consequence of many changes in agriculture and related industries, employers are seeking candidates with these types of growing skill sets, which are related to the current needs in agriculture [40].

In addition to identifying activities in graduate school that can assist students in improving their professional skills and consequently the ability to lead others, there were other graduate school experiences correlated with improved effectiveness of leadership in all professional roles. The three most significant experiences in graduate school for effective leaders were identified as ‘observing other leaders’, ‘job experiences’, and ‘mentoring students’. The majority of graduate students are starting their programs fresh out of undergraduate program, therefore lacking breadth in job experiences and presumably meaningful leadership experience.

Perhaps implementing internships, or other job responsibilities and expectations beyond student’s research could contribute to fill this gap. A quote from one of the respondents underscores this need: “As a result, too many of us learn on the job, making mistakes & re-inventing the wheel as we go (…)”. Observing other leaders is an ability that is not dissipated among young professionals, but it is present in everyday interactions. Our senses and observational skills can provide us with powerful information that helps us understand the people we interact with and their emotions, which in result can improve how we interact with people, and to detect skills that we need to develop [42]. Student mentorship can be difficult to achieve, mostly because effective mentoring relationships are not always available in academic settings and faculty lack formal training in mentoring [43]. Furthermore, graduate students in underrepresented groups often experience inadequate mentoring as a result of mentors’ inexperience in working with students in these groups [44].

Some skills recognized as crucial for effective leadership were also identified as top ranked among the survey respondents and included: ‘listen effectively’, ‘problem-solving’, ‘well-developed ethic, integrity and sense of loyalty’, ‘collaborative’, and ‘strategic and see the big picture’. Such skills can be adopted by university departmental units as indicative of offering professional skills training among graduate students that can help them to be more effective leaders during their graduate program, as well as outside of academia and in their professional careers. Training for those top ranked professional skills can be offered in many ways.

Our study has identified key recommendations that graduate programs should focus on to improve such skills for leadership development. Respondents from academia and industry unanimously recommended that graduate students need to have access to leadership opportunities and formal programs to increase leadership development. Opportunities were broadly mentioned as any experience that can increase leadership development among graduate students. But interestingly, formal programs were highly recommended among our respondents, which is in contrast to the aggregate data that showed that formal training, such as via courses, was not a significant experience for effective leadership (Fig. 3). Furthermore, most respondents (73.5%) indicated that such experiences were not available, which suggests that providing formal programs such as capstone courses, or part of the curriculum could be an efficient way to provide leadership training among graduate students in agricultural plant science disciplines. Moreover, some of these recommendations seemed to be dependent on employment type (academia vs. industry) of the participants. This was expected as each employment type demands different competencies. The following themes: real world experience, communication, teamwork, and business showed to be more relevant to industry. Few academic programs integrate real-world experience as part of learning and neither are there sufficient resources for faculty to experiment with how to refashion the way they teach or provide experiences that reflect the challenges that food and agriculture graduates will need in their future careers [40]. Communication and teamwork appear to be key for fruitful collaborations and to understand stakeholders, as this respondent said: “Effective oral and written communication are still highly valued skills in today’s marketplace, especially when interacting with customers/constituents (…) the more prepared a student is to demonstrate these skills, the more marketable they become in my opinion (…).” Furthermore, public speaking was also mentioned among industry respondents as an important recommendation for graduate student training.

The majority of professional and student participants reported in this study mentioned that they do not find graduate programs as a significant environment to prepare students for leadership roles in their future professional careers. Providing encouragement, professional training, and having graduate students mentoring undergraduates were the most common recommendations to improve leadership skills among academics. There are many benefits in mentoring other professionals as it can enhance productivity, increase engagement, and improve recruitment [45]. This recommendation is illustrated in the following quote from one of the respondents: “As I am employed by a Tier I research institution, the main emphasis is on research. The positive activities that seem to help graduate students develop leadership skills are (…) mentoring undergraduates in laboratories with the expectation of an end product (not just hands doing minor tasks).” Additionally, providing professional training, especially through participation in professional societies, offers a chance for the student to be exposed to their potential professional abilities in real-life situations when connecting with their colleagues or potential employers. The implementation for such recommendations is not trivial and should be aligned with the departmental or unit resources and student interest. Such alignment is challenging because the majority of departments in the agricultural plant sciences do not have the funding or faculty training necessary to adopt such measures.

The biggest barriers identified by the respondents of this survey to justify the lack of leadership development in the U.S. universities were because it is not part of the program curriculum and lack of advisor support. These barriers reflect on the importance for American universities to provide encouragement to departmental units, especially to advisors, for student leadership development, for example by providing career stimulus for faculty who successfully develop their graduate students professionally. Faculty can lack training in mentorship or even support from the university to provide such relationship, which can leave faculty unprepared for the duty of advising students [9]. Additionally, few granting agencies provide the financial support to faculty and students to extend professional training beyond research. The National Science Foundation is one of these few agencies that fund programs to prepare STEM graduate students to the workforce [2]. There are also efforts in a few universities that are adopting the splitting of Ph.D. programs to expand training opportunities that are not only focused on technical skills [39].

A theme that was surprisingly weak in our responses, were issues of Diversity, Equity, and Inclusion (DEI) in leadership training for graduate students. We assume that this is due, in part, as the last of our survey responses were collected just prior to the summer of 2020 when there was increased national attention on racial inequities. Even before 2020, the National Academy of Sciences, Engineering, and Medicine listed creating an environment and culture that promotes DEI in STEM graduate programs was one of the key issues of their cross-cutting themes in graduate education [5]. Among report recommendations was a focus on faculty development in DEI with the goal of improving mentorship skills. The quality of mentors-mentee relationships is an especially important factor in the development of scientific identity is underrepresented minority (URM) STEM students [46]. However, the mentorship needs of URM students is complex and factors like matching students with peer or near-peer identity mentors, the type of research experience, and opportunities to communicate their science identity to others should be tailored to individual students [47]. As agricultural plant science graduate students continue to come from more diverse backgrounds, issues involving DEI and the impacts on graduate student leadership development should be investigated.

In our study, a majority of student and professional participants did not believe graduate programs provided a significant contribution to their preparation for leadership roles in their future professional careers. This is may be a result of the perceived lack of leadership training in graduate programs, the non-leadership career pathways of some respondents, or that some professionals reported that post-graduate school experiences better filled gaps in leadership development. Regardless, these results point to a need to integrate leadership development and student training as the agricultural plant sciences field continue to evolve. Further, this study reinforces the need for academic program leaders and industry professionals to collaboratively develop meaningful leadership opportunities (e.g., internships, research team partnerships, leadership development workshops) for graduate students.

## Conclusions

Our survey identified that the majority of agricultural plant science professionals and students do not find graduate programs as an effective environment for providing leadership development, and that the lack of formal training, funding, and encouragement are significant barriers to move forward with such development in graduate programs. For leadership training, graduate students should participate in activities beyond research and teaching to remain competitive in the job search and increase leadership learning opportunities. For graduate programs, increasing formal programs and offering opportunities for leadership development are necessary for a successful generation of future leaders in both academic and industry employment settings. However, it is crucial to tailor such trainings with the employment type of choice. Our survey results also identified other experiences that can significantly contribute to the development of leaders, such as observing the behavior of other leaders, having job experiences beyond thesis / dissertation research, and effective mentorship during the student’s graduate program.

This survey was able to provide important findings that can help current and future graduate students and departmental units to focus on the current needs in leadership development and training opportunities. While we do not have all the answers to how to implement leadership development in graduate programs, we were able to identify experiences and relevant skills that are intertwined with relevant leadership training for future leaders in the agricultural plant science disciplines.

## Notes

### Competing Interest Statement

The authors have declared no competing interest.

